# Boosting transformation in wheat by BBM-WUS

**DOI:** 10.1101/2022.03.13.483388

**Authors:** Ziru Zhou, Yawen Yang, Guo Ai, Miaomiao Zhao, Baozhu Han, Chunjie Zhao, Yiqian Chen, Yuwei Zhang, Hong Pan, Caixia Lan, Qiang Li, Jieting Xu, Wenhao Yan

**Affiliations:** National Key Laboratory of Crop Genetic Improvement, Hubei Hongshan Laboratory, Huazhong Agricultural University, Wuhan, China; WIMI Biotechnology Co., Ltd, Changzhou, China

**Author notes:** **Correspondence**: Jieting Xu, Wenhao Yan.

**Keywords:** Agrobacterium-mediated transformation, BBM-WUS, GRF-GIF, wheat

## Abstract

Agrobacterium–mediated transformation is a cost-effective and convenient way to introduce foreign genetic elements to plants. However, only limited protocols successfully generates transgenic plants with fielder, a spring wheat variety. Wheat transformation method with higher efficiency and without genotype restriction is heavily demanded. Here, a heat-shock protocol, independent of Japan Tobacco had been established. Transgenic plants can be obtained from immature embryo within only 60 days by this protocol. Morphogenic regulators Baby boom and Wuschel (BBM-WUS) was proved to promote transformation efficiency for five to six times in wheat when co-infiltrated with agrobacterium containing target construct. Notably, half of the transformants are BBM-WUS free and moreover, the BBM-WUS containing plants could be picked by florescent marker that was co-expressed with BBM-WUS. In conclusion, we managed to establish a new wheat transformation protocol with shorter duration than published protocol.

## Results

Transgenic technology greatly promotes function genomic study and precision breeding in crops. Compared with biolistic bombardment, agrobacterium-mediated transformation is a low-cost and more favorable method in terms of clearer and easier genetic lineage of the transgenic lines with very few integrated T-DNA copies. Till now, agrobacterium-mediated transformation methods have been established in all three major crops, including rice, maize and wheat. Rice transformation system uses callus induced from mature seeds but only immature embryo could be used to perform transformation in maize and wheat (Hiei and Komari, 2008; Huw and Peter, 2009; Sidorov and Duncan, 2009). A morphological gene pair, Baby boom and Wuschel (BBM-WUS) has been used to boost transformation in maize. BBM-WUS induces somatic embryo to facilitate transformation shortly after infiltration (Lowe et al., 2016). In wheat, Japan Tobacco Company developed a protocol to perform agrobacterium-mediated transformation (Ishida et al., 2015). Based on this method, Debernardi and colleagues discovered the morphological gene, GROWTH-REGULATING FACTOR 4 and its cofactor GRF-INTERACTING FACTOR 1 (GRF-GIF) chimera as a booster for wheat transformation, which was later been verified in a transient expression assay (Debernardi et al., 2020; Qiu et al., 2021). However, the method developed by Japan Tobacco has been patented and is not fully open, so only a few organizations that paid the fee could use the method with restrictions. Development of new protocols for efficient agro-transformation in wheat is highly demanded. Except for GRF-GIF, whether BBM-WUS would promote wheat transformation remains unknown.

In order to establish the agrobacterium-mediated transformation method in wheat, we first tried the methods published by JP and another study from England (Hayta et al., 2019). Unfortunately, we hardly obtain any transformants from either method, possibly due to the failure of infection, which was indicate by nonexistent GFP signal and super unhealthy immature embryo after selection. We then tested different combinations among parameters of medium-components, embryo size and infiltration method to figure out an optical combination. We found that a heat-shock protocol worked efficiently. In this protocol, we incubated bacterium with immature embryo at 42 degree for 2 minutes. The best immature embryo size for transformation is 1.5-2.0mm and our self-developed co-culture medium worked best. Strong GFP signal could be observed after co-inoculation. The infected embryo was transferred to resting medium after co-cultivation. One week later, the embryo was submitted to selection medium. Notably, we combined selection and regeneration as one single step which largely shortens the whole process to around 40 days before we could proceed rooting. Following the method we described above, we successfully obtain transgenic plants from immature embryo within 60 days with an average value of 3.72% (34/902) (Figure 1a).

**Figure 1.**
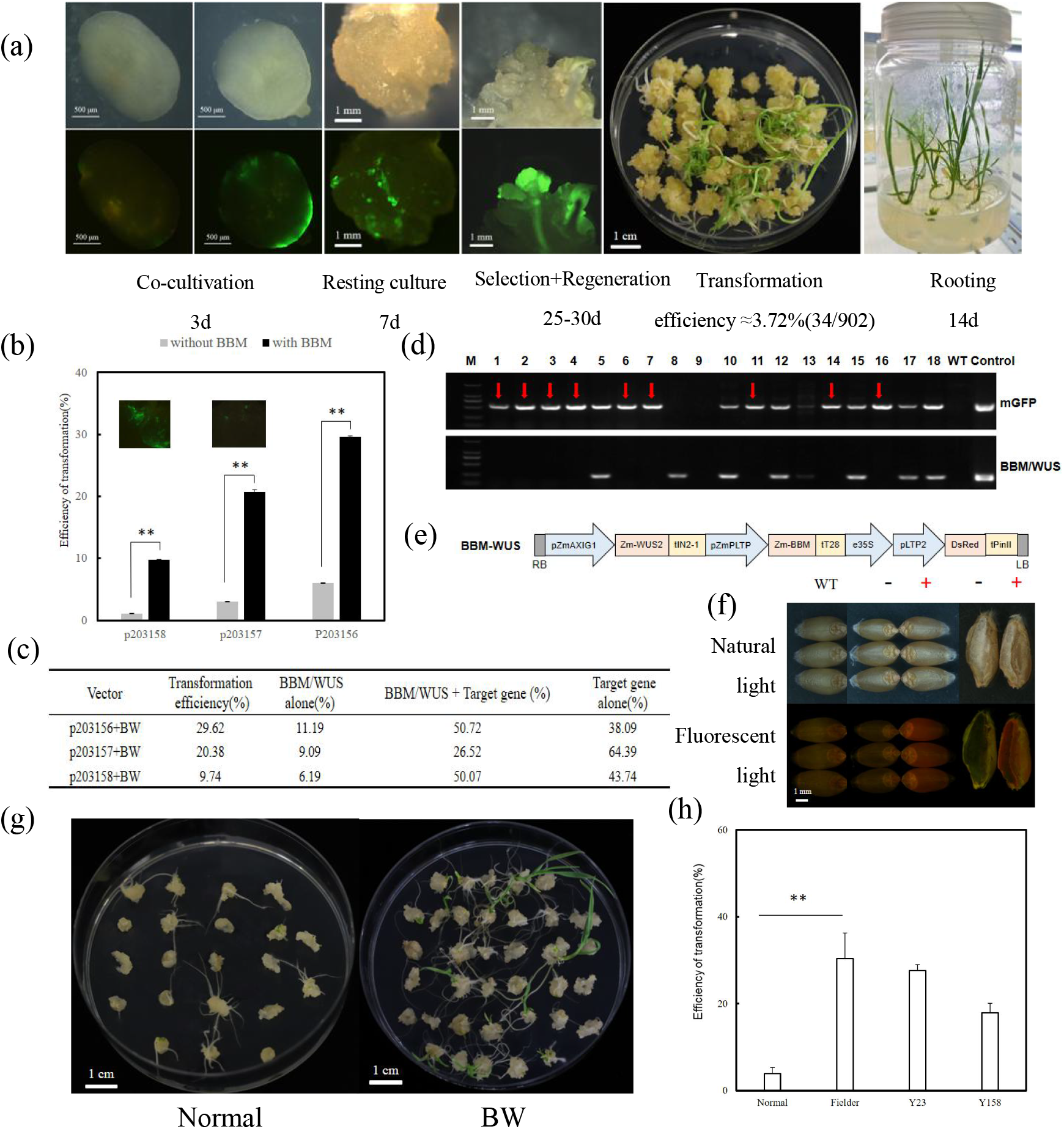
Efficient agrobacterium-mediated transformation methods in wheat boosted by BBM-WUS and GRF-GIF. (a). Complete duration from immature embryo to transgenic plants.(b-d) BBM-WUS significantly increased transformation efficiency and half are BBM-WUS free transgenic plants.p203156 is an empty vector;p203157 contains mGFP and p203158 contains enhanced GFP (eGFP). The red arrow in the DNA gel picture indicates BBM-WUS free transformants and the table shows the presentence of BBM-WUS and targeted DNA. (e) Schematic diagram of BBM-WUS construct containing red florescent marker. (f) Visualization of red florescent signal. (g) Performance of infected embryo with or without booster at regeneration stage. (h) Transformation efficiency in Yangmai23 (Y23) and Yangmai158 (Y158) by using BBM-WUS.

Inspired by the fact that morphological genes could enhance transformation, we fist tested whether BBM-WUS could accelerate transformation in wheat. BBM-WUS induces neighboring cells to form somatic embryo to be easily infected by agrobacterium thus eliminate the T-DNA of morphological genes. Strikingly, BBM-WUS elevated the positive rate from 1.14% to 9.74% and 3.03% to 20.38% when co-infiltrated with constructs which possess strong eGFP and moderate mGFP signal, respectively. For the empty vector, the positive rate was increased from 6.03% to 29.62% (Figure 1b). The expression level of GFP reporter affects transformation efficiency here. In general, BBM-WUS helps to enhance transformation efficiency for five to six times than the one without morphological gene. As expected, of all transformants, around half of the positive transgenic plants contain only GFP but not BBM-WUS thus to eliminate the growth defects caused by BBM-WUS (Figure 1c,d). Moreover, the aleurone specially expressed red florescence marker was placed together with BBM-WUS within one T-DNA to facilitate removing seeds expressing BBM-WUS(Xu et al., 2021)(Figure 1e). Although the red color is not easy to be recognized by naked eyes, the seeds containing the red florescent marker can be clearly distinguished from the ones without the marker under fluorescent light (Figure 1f).

One big challenge of bacterium-mediated transformation is that the efficiency is tightly linked with genotype. Currently, only a few accessions including fielder, bobwhite and Kenong199 are reported to be easily transformed. Manipulating elite wheat is more desirable for wheat improvement. Yangmai23 and Yangmai158, two elite varieties that are widely grown along Yangzi River were used to test whether our BW method could expand the genotypes. We successfully got positive transformants from both accessions (Figure 1h). This result convinced us that the newly developed agrobacterium-mediated wheat transformation method plus morphological genes is an efficient solution to break the genotype restriction in wheat.

In conclusion, we managed to establish an efficient wheat transformation protocol independent of Japan Tobacco protocol. In this protocol, morphological genes BBM-WUS were found to boost agrobacterium-mediated transformation in wheat for the first time. Future effect should be investigated to identify novel boosters, such as downstream causal targets of BBM-WUS.

## Author Contributions

Z.Z., Y.Y., C.Z. and B.H. tested the conditions and established the protocol. Z.Z., Y.C., and H.P. conducted the genotyping and collected the data. M.Z., Y.Z., C.L., and Q.L. provided the immature embryos. J.X. generated the constructs with help from Z.Z and G.A. W.Y. analyzed the data and wrote the manuscript with input from J.X. J.X and W.Y. conceived of the study. All authors read and approved the final manuscript.

## Acknowledgements

This work was supported by the National Key Research and Development Program of China (2020YFE0202300) and the Fundamental Research Funds for the Central Universities (2662020ZKPY002).

## References

1. Debernardi J M, Tricoli D M, Ercoli M F et al. A GRF-GIF chimeric protein improves the regeneration efficiency of transgenic plants. Nature Biotechnology, 2020, 38:1274–1279

2. Gordon-Kamm B, Sardesai N, Arling M et al. Using Morphogenic Genes to Improve Recovery and Regeneration of Transgenic Plants. Plants, 2019, 8,1–15

3. Hayta S, Smedley M A, Demir S U et al. An efficient and reproducible Agrobacterium-mediated transformation method for hexaploid wheat (Triticum aestivum L.). Plant Methods, 2019, 15:121

4. Hiei Y and Komari T. Agrobacterium-mediated transformation of rice using immature embryos or calli induced from mature seed. Nat Protocol, 2008, 3:824–834

5. Huw D and Peter R. Transgenic Wheat, Barley and Oats. Methods In Molecular Biology, 2009, 3–20

6. Ishida Y, Tsunashima M, Hiei Y et al. Wheat (Triticum aestivum L.) transformation using immature embryos. Methods In Molecular Biology, 2015, 1223:189–198

7. Lowe K, Wu E, Wang N et al. Morphogenic Regulators Baby boom and Wuschel Improve Monocot Transformation. The Plant Cell, 2016, 28:1998–2015

8. Qiu F, Xing S, Xue C et al. Transient expression of a TaGRF4-TaGIF1 complex stimulates wheat regeneration and improves genome editing. Sci China Life Science, 2021, doi: 10.1007/s11427-021-1949-9

9. Xu J, Yin Y, Jian L et al. Seeing is believing: a visualization toolbox to enhance selection efficiency in maize genome editing. Plant Biotechnology Journal, 2021, 19:872–874

